# SGLT2 inhibitors activate pantothenate kinase in the human heart

**DOI:** 10.1101/2024.07.26.605401

**Authors:** Nicholas Forelli, Deborah Eaton, Jiten Patel, Caitlyn E. Bowman, Ryo Kawakami, Ivan A. Kuznetsov, Kristina Li, Claire Brady, Kenneth Bedi, Yijun Yang, Kaustubh Koya, Emily Megill, Daniel S. Kanter, Louis G. Smith, Gregory R. Bowman, Nathaniel Snyder, Jonathan Edwards, Kenneth Margulies, Zoltan Arany

## Abstract

Inhibitors of sodium glucose cotransporter-2 (SGLT2i) demonstrate strong symptomatic and mortality benefits in the treatment of heart failure but appear to do so independently of SGLT2. The relevant pharmacologic target of SGLT2i remains unclear. We show here that SGLT2i directly activate pantothenate kinase 1 (PANK1), the rate-limiting enzyme that initiates the conversion of pantothenate (vitamin B5) to coenzyme-A (CoA), an obligate co-factor for all major pathways of fuel use in the heart. Using stable-isotope infusion studies, we show that SGLT2i promote pantothenate consumption, activate CoA synthesis, rescue decreased levels of CoA in human failing hearts, and broadly stimulate fuel use in *ex vivo* perfused human cardiac blocks from patients with heart failure. Furthermore, we show that SGLT2i bind to PANK1 directly at physiological concentrations and promote PANK1 enzymatic activity in assays with purified components. Novel *in silico* dynamic modeling identified the site of SGLT2i binding on PANK1 and indicated a mechanism of activation involving prevention of allosteric inhibition of PANK1 by acyl-CoA species. Finally, we show that inhibition of PANK1 prevents SGLT2i-mediated increased contractility of isolated adult human cardiomyocytes. In summary, we demonstrate robust and specific off-target activation of PANK1 by SGLT2i, promoting CoA synthesis and efficient fuel use in human hearts, providing a likely explanation for the remarkable clinical benefits of SGLT2i.

## Introduction

Sodium glucose cotransporter-2 inhibitors (SGLT2i) are a class of oral-antihyperglycemics that have surprisingly also become a foundational part of guideline directed medical therapy (GDMT) for patients with heart failure (HF)^1^. The drugs were originally developed to promote urinary loss of glucose in patients with type 2 diabetes mellitus (T2DM) by inhibiting SGLT2-mediated reabsorption of glucose in the renal proximal tubules. In phase III trials in patients with T2DM, SGLT2i reduced major adverse cardiovascular events, but also demonstrated a strong protection against HF, independently of T2DM severity^2^. Subsequent trials revealed that treatment with SGLT2i reduce rates of death and heart failure rehospitalization equivalently in HF patients with or without diabetes^3-7^. SGLT2i are now recommended as first-line agents in the treatment of HF. However, the side effects of SGLT2i are non-negligible, including the on-target side effect of genital mycotic and urinary tract infections, presumed to be caused by the high levels of glucose in the urine. Additionally, SGLT2i and other oral anti-hyperglycemics are often halted upon hospital admission in preference of using insulin for glycemic control, thereby potentially mitigating their cardiovascular benefit in patients hospitalized with HF exacerbations.

The mechanism by which SGLT2i achieve cardioprotection and these profound clinical benefits remains unclear. An off-target mechanism of action is highly likely because: (1) SGLT2 is not expressed in the heart, but several studies have shown direct effects of SGLT2i on isolated cardiomyocytes or hearts^8^, and (2) treatment with SGLT2i in mice reduces infarct size after ischemia/reperfusion^9^ and partially improves function after trans-aortic constriction/myocardial infarction equivalently in animals genetically lacking SGLT2 and control animals^10^. The identity of this non-SGLT2 pharmacological target of SGLT2i remains elusive. Here we show that SGLT2i have a direct and profound effect on metabolism and contractile function in human hearts, and we identify the enzyme pantothenate kinase (PANK1) as a novel target of SGLT2i that explains these benefits.

### SGLT2i directly promote human cardiac metabolism

There is ongoing debate on whether SGLT2i benefit HF by directly affecting the heart or via broader systemic metabolic reprogramming, but data in humans are lacking.^11-13^ We therefore first sought to test if SGLT2i directly affect metabolism in human myocardium. We developed a method to perfuse large cardiac tissue blocks *ex vivo* (average >40g), taken from the interventricular septum (IVS) of either failing hearts of transplant recipients or non-failing hearts from organ donors (see Methods). Tissue blocks were perfused with a recirculating Krebs-Henseleit buffer containing cardiac fuels at physiological concentrations, several of which were labeled with stable heavy isotopes ([6,6-^2^H]-glucose, [1-^13^C]-glutamine, [3-^13^C]-lactate, [U-^13^C]-valine, and [U-^13^C]-3-hydroxybutyrate) and then treated with either 700nM of the SGLT2i empagliflozin (EMPA) or vehicle control (Figure 1A). The chosen EMPA dose reflects plasma levels achieved in treated patients^14-16^. Multiple IVS blocks from each heart were cannulated and perfused in parallel such that every heart served as its own control. We first tested the impact of EMPA on cardiac fuel consumption by measuring the disappearance of labeled nutrients over time in the recirculating perfusate. Strikingly, treatment with EMPA increased the uptake of all fuels in failing human myocardium, indicating a broad activation of metabolic activity by EMPA (Figure 1B). Consistent with this, the tissue levels of these fuels were significantly elevated in the cardiac blocks treated with EMPA (Figure 1C).

**Figure 1:**
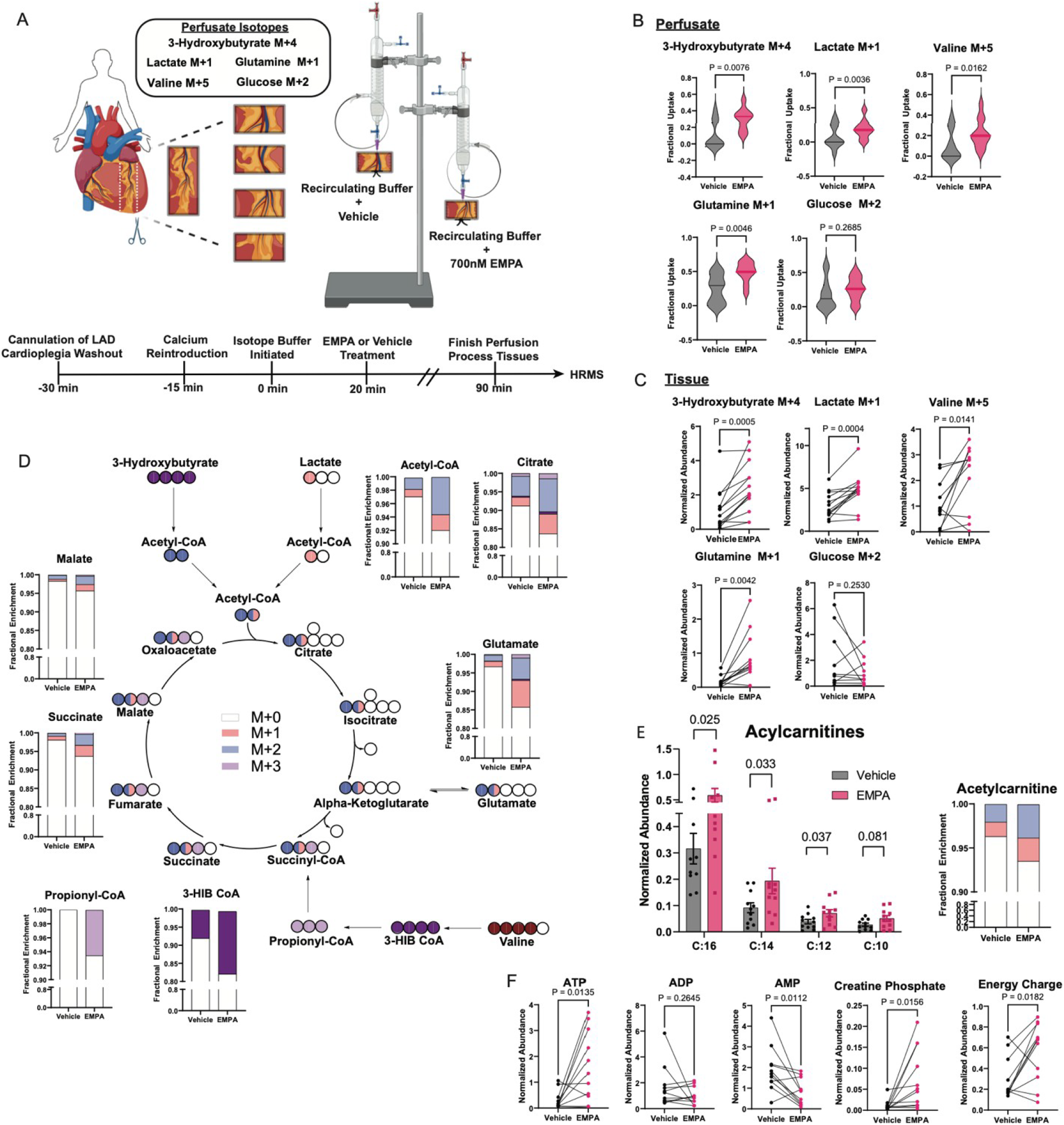
SGLT2i directly promote human cardiac metabolism. **A**, experimental design. **B**, Fractional uptake of indicated substrates from the perfusate after 90 minutes. **C**, Normalized abundance of indicated substrates after 90 minutes of perfusion. **D**, Fractional ^13^C enrichment of the indicated intermediate metabolite after 90 minutes of perfusion. **E-F**, Relative tissue levels of indicated metabolites after 90 minutes of perfusion. P-values by paired 2-sided student’s t-test. Error bars are ±SE.

The presence of isotopically labeled fuels in the perfusate allowed us to next trace the incorporation of fuel-derived carbons into the tricarboxylic acid cycle (TCA), an indirect measure of fuel oxidation. The fractional enrichment of labeled TCA carbons was markedly increased in cardiac blocks treated with EMPA (Figure 1D), reflecting increased incorporation of labeled carbons from lactate and 3-hydroxybutyrate via acetyl-CoA, and from valine via propionyl-CoA and succinyl-CoA. Although the perfusate contained no labeled fatty acids, elevated acyl-carnitines in EMPA-treated cardiac blocks suggested that fatty acid oxidation was also increased by EMPA treatment (Figure 1E). Finally, consistent with a broad activation of metabolic activity, EMPA treatment substantially increased ATP and decreased AMP, thus boosting the energy charge in these cardiac tissues (Figure 1F). We conclude that EMPA directly and broadly promotes oxidative metabolism in human hearts, thereby rescuing the well-established energetic defect seen in failing hearts^17-19^.

### SGLT2i activate CoA synthesis to promote cardiac metabolism

To begin to understand how SGLT2i so broadly affect cardiac metabolism, we performed untargeted global metabolomics on perfused heart blocks treated with EMPA versus vehicle control (Figure 2A). Among several metabolites significantly affected by EMPA treatment, we noted the more than 2-fold depletion of pantothenate (Figure 2A). Pantothenate, also known as vitamin B5, is the precursor to the synthesis of coenzyme A (CoA; Figure 2B right panel), which is an obligate co-factor for nearly all pathways of cellular fuel use, including degradation of carbohydrates, fats, and amino acids^20^. EMPA significantly increased the ratio of phospho-pantothenate to pantothenate in the perfused cardiac blocks (Figure 2A, right panel), consistent with activation of the CoA biosynthetic pathway. To formally test the consumption of pantothenate, we supplemented the perfusate with [^13^C_3_15N]-pantothenate. Treatment with EMPA more than doubled the fractional uptake of pantothenate by the human hearts, and markedly increased incorporation of isotopically labeled pantothenate into acetyl CoA (AcCoA) (Figure 2B, left panels). Thus, the data suggest the possibility that EMPA promotes CoA synthesis, rescuing insufficient CoA abundance in failing hearts, and thereby explaining the broad salutary effects observed with EMPA treatment. Consistent with this notion, the abundance of various acyl-CoAs, and most importantly free CoASH, are markedly decreased in human failing hearts, compared to nonfailing donors (Figure 2C).

**Figure 2:**
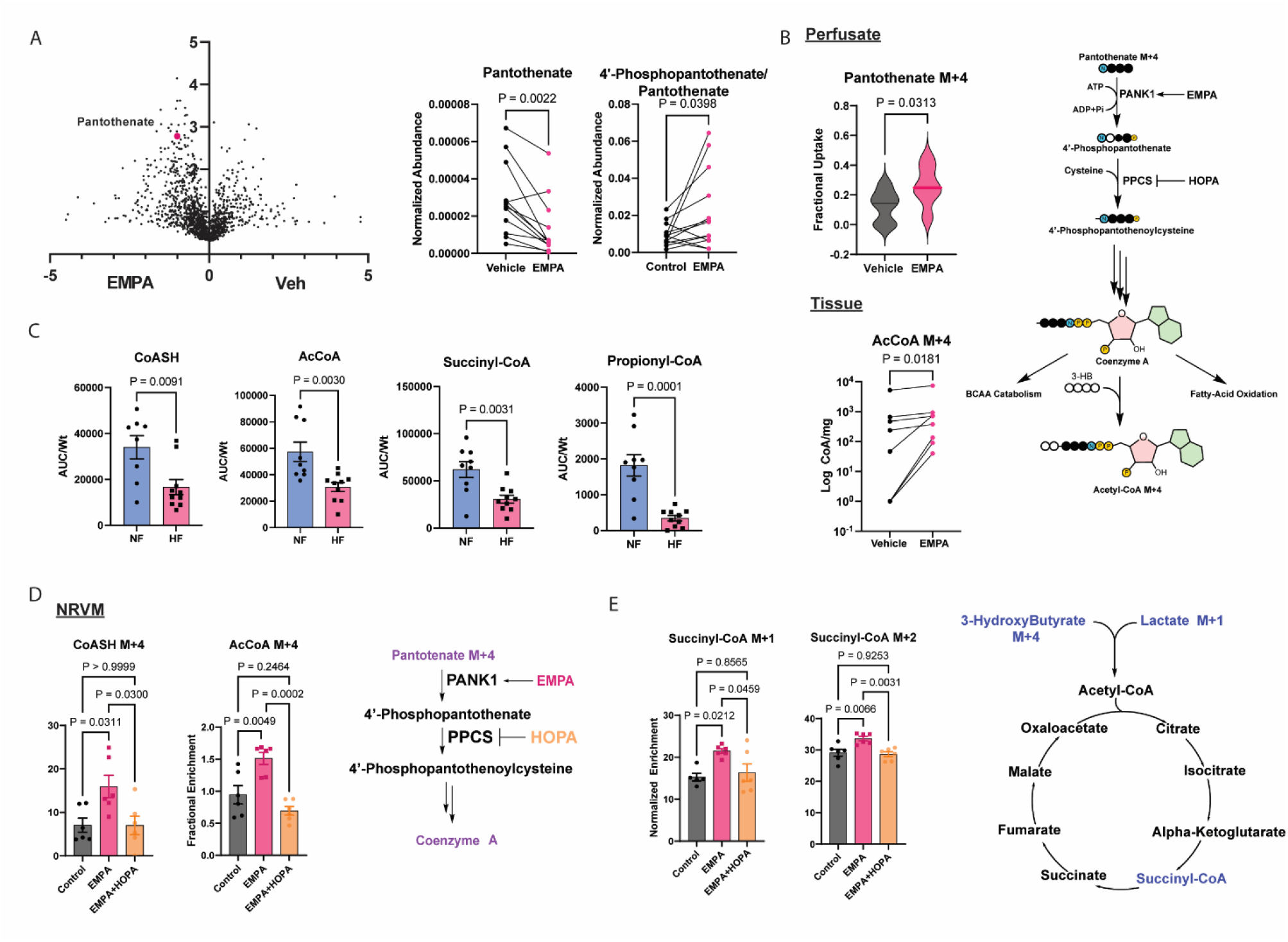
SGLT2i activate CoA synthesis to promote cardiac metabolism. **A**, Left: Volcano plot of fold-change in perfused human cardiac tissue metabolite abundance with EMPA treatment. Middle: depletion of pantothenate. Right: increase in phospho-pantothenate/pantothenate ratio. **B**, Top left: increase in fractional uptake of pantothenate by perfused human cardiac blocks in response to EMPA treatment. Right: diagram of CoA synthesis from pantothenate. Bottom left: increased labeling of AcCoA from labeled pantothenate precursor, in the presence or absence of EMPA treatment. **C**, Decreased abundance of CoA species in post-transplant human heart failure samples, compared to donor controls. **D-E** Fractional enrichment of AcCoA from labeled pantothenate (**D**) and succinyl-CoA from 13C-labeled cocktail (**E**) in NRVMs treated with vehicle vs EMPA vs EMPA plus the PPCS inhibitor HOPA. P-values by paired 2-sided student’s t-test, except **c** which is unpaired. Error bars are ±SE.

To test if the activation of CoA synthesis by SGLT2i is required for the effects of SGLT2i on the TCA noted above (Figure 1), we turned to cultured cardiomyocytes and the use of hopantenate (HOPA), which blocks CoA synthesis by inhibiting phosphopantothenoylcysteine synthase (PPCS), the second enzyme in the CoA synthesis pathway (Figure 2B). As with human blocks, treatment of neonatal rat ventricular myocytes (NRVMs) with EMPA significantly increased the incorporation of isotopically labeled pantothenate into AcCoA as well as free CoASH, and this process was prevented by the addition of HOPA (Figure 2D). Also as seen in the human cardiac blocks, treating NRVMs with EMPA promoted the incorporation of fuel-derived carbons into the TCA cycle (Figure 2E), demonstrating that the effects of EMPA on cardiomyocytes are direct, independent of blood flow, the vasculature, or other cells. Importantly, the addition of HOPA prevented the EMPA-mediated boost in TCA labeling (Figure 2E). We conclude that the activation of CoA synthesis by EMPA is required for the boosting effects of EMPA on the TCA.

### SLGT2i activate PANK1

We next sought to identify the enzyme target of SGLT2i that mediates the observed boost in CoA synthesis. The five-step process of CoA synthesis from pantothenate (Figure 2B) begins with the rate-limiting phosphorylation of pantothenate by pantothenate kinase (PANK). Three genes encode PANKs, of which PANK1 is most expressed in cardiomyocytes^20^. Interestingly, PANK1 expression is suppressed in failing human hearts, and pantothenate levels tend to be increased (Figure 3A). We therefore hypothesized that SGLT2i activate PANK1 to promote CoA synthesis. Consistent with this notion, EMPA significantly increased the ratio of phospho-pantothenate to pantothenate in perfused cardiac blocks (Figure 2A). To test directly if EMPA reaches and binds to PANK1, we first used a cellular thermal shift assay^21^ (CETSA). HEK293 or HEPG2 cells, which express PANK1, were treated with EMPA versus vehicle control and then subjected to incremental increases in ambient temperature, followed by western blotting for PANK1. The addition of EMPA strongly increased the thermal stability of PANK1 (Figure 3B and Supplementary Figure 1A), similar to that achieved by a previously described pan-PANK activator (PZ-2891^22^; Supplementary Figure 1B), demonstrating efficient entry of EMPA into cells and ligand-induced thermal stabilization. To test for direct binding of EMPA to PANK1, we immobilized EMPA on beads and precipitated associated proteins from HEK293 cellular extracts. EMPA-loaded beads, but not control beads, efficiently bound PANK1 from these extracts (Figure 3C), demonstrating direct binding. In an orthogonal approach, immunoprecipitation of PANK1 from HEK293 cells also co-precipitated EMPA, as detected by mass spectrometry (Figure 3D). We conclude that EMPA binds to PANK1.

**Figure 3:**
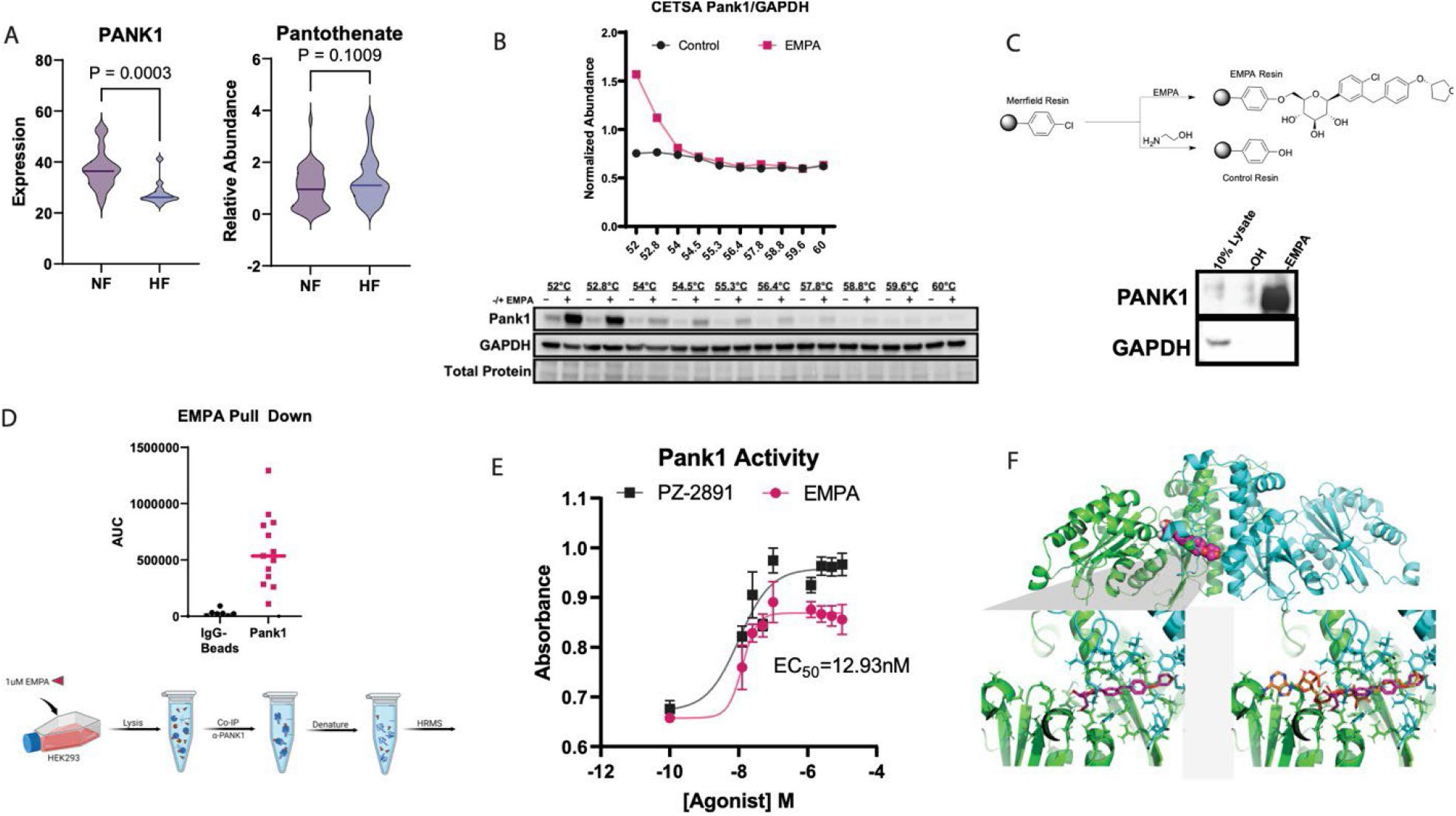
SGLT2i activate PANK1. **A**, mRNA levels of *PANK1* expression (left) and levels of pantothenate (right) in human failing and nonfailing hearts. **B**, CETSA assay of PANK1. **C**, Binding of PANK1 to EMPA-immobilized beads. **D**, Co-immunoprecipitation of EMPA with PANK1. **E**, PANK1 enzymatic activity in response to increasing doses of known PANK1 activator (PZ-2891) and to EMPA. **F**, *in silico* modeling shows EMPA binding in same allosteric pocket within PANK1 as bound by acetyl-CoA. Top: ribbon model of PANK1 dimer. Bottom left: zoom-in of pantothenate binding to PANK1. Bottom right: super-imposed binding of AcCoA, showing extension into the PANK1 enzymatic pocket. P-values by unpaired 2-sided student’s t-test. Error bars are ±SE.

We next tested the direct effects of EMPA on PANK1 enzymatic activity, using PANK1 enzyme generated and purified from bacterial expression and two different kinase activity assays^23,24^, including one previously established to quantify PANK enzymatic activity^24^. Treatment of purified PANK1 with EMPA activated pantothenate kinase activity to a similar extent as a previously described pan-PANK activator (PZ-2891^22^) (Figure 3E and Supplementary Figure 2). Activation was achieved with a EC50 of 13 nM (Figure 3E), well below plasma concentrations achieved with EMPA therapy in humans (peak 500-1500 nM^14-16^), and comparable to PZ-2891. We conclude that EMPA, at physiologically relevant concentrations, binds to and activates enzymatic activity of PANK1.

To begin to understand how SGLT2i activate PANK1, we leveraged PopShift, a novel *in silico* modeling methods^25^. We used this approach to analyze what poses for EMPA and acyl-CoA were compatible with the receptor conformations sampled from 5 milliseconds of aggregate molecular dynamics of the unliganded PANK1 homodimer in explicit solvent, which we collected with Folding@home. The poses we collected predicted that EMPA binds to PANK1 within an allosteric pocket that can also be occupied by acyl-CoA species (Figure 3F). Acylated CoA species inhibit PANK1, in a negative feedback loop, by occupying this pocket and reaching into the enzymatic active site, thereby sterically inhibiting enzymatic activity^26^. Importantly, when EMPA occupies this allosteric pocket, it does not reach into the enzymatic active site and is thus not predicted to hinder enzymatic activity. We therefore predict that EMPA activates PANK1, at least in part, by displacement of inhibitory acyl-CoA species. Other allosteric mechanisms may also contribute.

### PANK1 activation by SGLT2i is required for functional benefits of SGLT2i

SGLT2i have profound clinical benefits in patients with HF, and we show above that SGLT2i directly improve human cardiac metabolism. To test if SGLT2i also directly improve human cardiomyocyte function, we isolated adult human cardiomyocytes from failing and non-failing donor hearts and quantified measures of contractility and relaxation, as previously described^27^ (Figure 4A). The addition of EMPA to the cells significantly increased the fractional shortening and peak height of contraction, markers of contractility (Figure 4B-C). EMPA also significantly increased relaxation velocity and decreased time to return to 90% baseline, indicating improvements in relaxation, an energetically demanding process that requires the rapid reuptake of calcium into the sarcoplasmic reticulum (Figure 4B-C and Supplementary Figure 3). We conclude that SGLT2i improve human cardiomyocyte contractility and relaxation directly, independently of systemic effects, hormonal context, or input by the vasculature.

**Figure 4:**
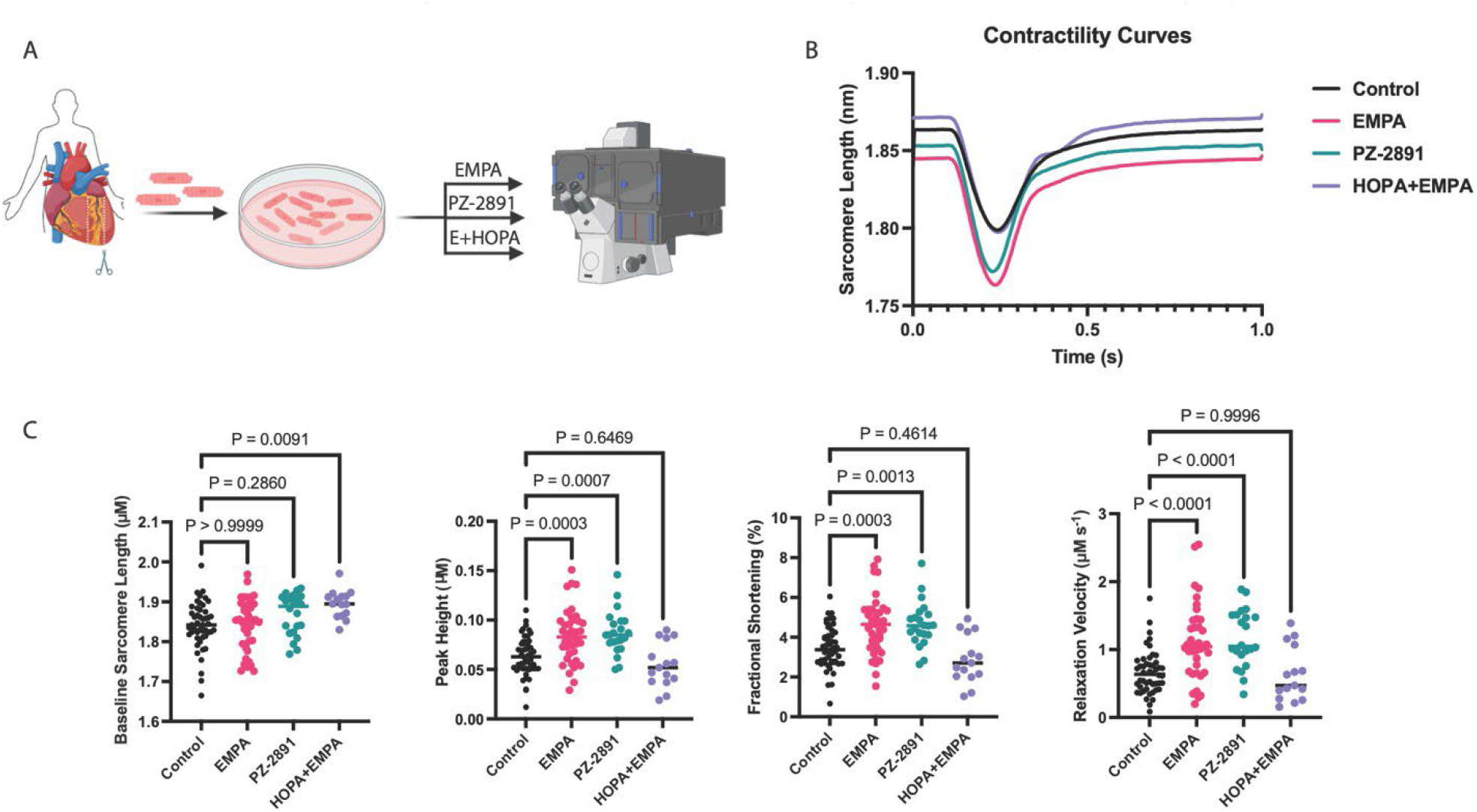
PANK1 activation by SGLT2i is required for functional benefits of SGLT2i. **A**, experimental outline. **B-C**, Sample contractility and relaxation curves (**B**) and quantification of parameters (**C**) from single human adult cardiomyocytes treated with the indicated agents. P-values are by one-way ANOVA.

We next tested the role of PANK1 activation by EMPA in its effects on cardiomyocyte contractile function. The addition of PZ-2891 to the human cardiomyocytes precisely mimicked the effects of EMPA on peak height, fractional shortening, and relaxation velocity (Figure 4B-C), demonstrating that PANK1 activation is sufficient to promote cardiomyocyte function.

Conversely, the addition of HOPA to the EMPA-treated cells completely reversed the beneficial effects of EMPA (Figure 4B-C). We conclude that PANK1 activation by SGLT2i is both necessary and sufficient for the functional improvements in contractility and relaxation conferred by SGLT2i on human cardiomyocytes.

## Discussion

We demonstrate here a critical off-target effect of SGLT2i: the direct activation of PANK1, resulting in the acute stimulation of CoA synthesis. We also demonstrate that activation of PANK1 by SGLT2i directly promotes human cardiac metabolism and contractility, independently of kidney function, metabolic and hormonal milieu, or input from the vasculature. These observations thus provide a potential explanation for the remarkable clinical benefits achieved by treating HF patients with SGLT2i, a benefit unlikely to be mediated by inhibition of SGLT2 itself because SGLT2 is not expressed in the heart, and because the benefits are also seen in mice genetically lacking SGLT2^9,10^. Activation of PANK1 by SGLT2i may also explain the surprisingly rapid benefits of SGLT2i, often seen within days of treatment initiation. The identification of PANK1 as a relevant off-target of SGLT2i opens the possibility of developing novel agents that more potently or specifically target PANK1, potentially both increasing efficacy and avoiding the on-target side effects of SGLT2i, euglycemic diabetic ketoacidosis, and UTIs.

Our work strongly implicates CoA biology in HF pathogenesis. CoA synthesis is central to oxidative metabolism, on which the heart primarily depends. We and others have shown here and elsewhere^28^ that both free CoA and esterified CoA species are strongly reduced in human failing hearts, suggesting that CoA abundance may be limiting. Our findings here that PANK1 activation with SGLT2i or a known PANK activator promote human cardiomyocyte metabolism and contractility strongly support this notion. Mice lacking cardiac PANK1 develop cardiomyopathy in response to hemodynamic challenges^29^, underscoring the critical role of PANK1 in cardiac function. Similarly, homozygous or compound heterozygous mutations in PPCS, the second enzyme in the CoA biosynthesis pathway, cause DCM in humans^30^, demonstrating the key role of CoA synthesis in human cardiac function.

The systemic benefits of SGLT2i may also reflect actions on PANK1 outside the heart. In addition to cardiomyocytes, PANK1 is also expressed in hepatocytes, renal tubular cells, gut epithelium, and neurons^31^. Interestingly, SGLT2i have shown benefits in NAFLD and CKD, likely independent of effects on cardiac function^32,33^. Activation of PANK1 in these tissues may thus in part explain these clinical benefits of SGLT2i. SGLT2i may also modulate other isoforms of PANK. PANK1 is the dominant PANK isoform in the heart, accounting for >70% of pantothenate activity, but both PANK2 and 3 are also present and activating these other isoforms may further extend the extracardiac benefits of SGLT2i.

In summary, we identify PANK1 as a direct target of SGLT2i, and demonstrate that stimulation of PANK1 and CoA synthesis by SGLT2i improves human cardiac metabolic and contractile activities. The results likely at least in part explain the remarkable benefits of SGLT2i in patients with HF and provide the opportunity to design novel compounds with potentially higher efficacy and fewer SGLT2-mediated side effects.

## Methods

### Human Heart Procurement

Procurement of human myocardial tissue was performed under protocols and ethical regulations approved by the Institutional Review Board at the University of Pennsylvania and the Gift-of-Life Program (Pennsylvania) as previously described.^34,35^ Prospective informed consent for research use of explanted heart tissue was obtained from transplant recipients in the case of failing hearts and from next-of-kin in the case of hearts from deceased organ donors. *In vivo* function was assessed by echocardiography of all hearts. At the time of procurement, hearts were arrested *in situ* with ice-cold cardioplegia and transported to the laboratory on wet-ice with cardioplegia within two hours.

### Preparation of Human Myocardial Blocks for Perfusion

Upon receipt, the interventricular septum (IVS) was dissected from the right and left ventricle taking care to preserve the architecture of the left anterior descending (LAD) artery. After separation of the IVS with intact LAD, segments of cardiac tissue were divided such that multiple portions of IVS had an intact segment of LAD and septal perforators for cannulation and perfusion. The LAD segments were cannulated with a 16-20 gauge cannula, and secured in place by ligation with suture or surgi-clip. The distal end of the perfusing LAD segment, as well as large branching vessels were ligated to allow for perfusion through the myocardium.

### Stable Isotope Perfusion of Human Myocardial Blocks

The cannulated human myocardium was attached to a langendorff apparatus and perfused with Krebs-Henseleit buffer solution (KHB) for 15 minutes to allow for rewarming and removal of cardioplegia. The KHB was continually oxygenated. After 15 minutes, calcium was introduced stepwise to a final concentration of 1mM in the KHB. After calcium reintroduction, the buffer was swapped to a modified KHB containing the stable-isotope metabolites (5mM 6,6-D2 Glucose (50%), 1.2mM 3-13C Lactate (50%), 0.4mM 13-C5 Valine (50%), 0.3mM 13C-4 D-3-hydroxybutyrate (100%), 0.5mM 1-13C glutamine (100%), with or without 0.002mM 3-13C-1-N15 Pantothenate (100%)). Samplings of the perfusate were taken at time 0, and every 10-minutes thereafter. After 20-minutes of perfusion with the isotope buffer, myocardial blocks were treated with vehicle (DMSO, 0.0008% final solution) or empagliflozin (700nM final concentration) by addition to the recirculating buffer. The experiment was allowed to proceed for a total of 90-minutes, at which time the myocardial blocks were cut down and rapidly freeze-clamped in liquid nitrogen.

### Human Cardiomyocyte Contractility

Myocytes were isolated as described previously.^34,35^ Briefly, the IVS was dissected, and the LAD architecture was maintained. The LAD was cannulated with an 16-20 g cannula and perfused in a Langendorff incubation chamber, maintaining a temperature of 37°C. The IVS tissue was perfused with Ca^2+^ free solution (KHB containing 20 mM 2,3-butanedione monoxime (BDM),10 mM taurine, 0.3mM Valine, 0.3mM Leucine, 0.3mM Isoleucine, 0.3mM 3-hydroxygutyrate) for 5 minutes. Then, 200mL of KHB containing 294 U/mL of collagenase was perfused for 25 minutes. Ca^2+^ was reintroduced over the course of 15 minutes. Finally, the tissues were perfused for 5-minutes with KHB containing 1% BSA. The myocardium was removed from the cannula and minced. The resulting cell-suspension was filtered through a 280-um nylon mesh (Component Supply U-CMN-289), and resuspended in M-199 containing 10% FBS, 1% Primocin, and 0.1% cytochalasin.

### Cardiomyocyte Contractility

Cells were cultured overnight at 37°C with 5% CO_2_ in uncoated 6-well dishes. Cells were treated 90-minutes before measure contractility with either: vehicle (DMSO, 0.01%) empagliflozin (700nM, or 100uM), PZ-2891 (700uM or 100uM), calcium hopantenate (25uM), or empagliflozin+hopantenate (700nM+25uM, respectively) in M-199 medium. Contractility measurements were performed as described previously.^34,35^ Briefly, cardiomyocytes were subjected to a field stimulation at of 20V at 1Hz with a myopacer (IonOptic MYP100).

Sarcomere length changes were measured after 10-15 seconds of 1 Hz pacing to achieve steady state by a high-speed video image acquisition Nikon PU-2000 inverted confocal microscope. Fourier transform and analysis of contractility data was performed (Ionwizard, IonOptix). A minimum of 5 steady state contractions for each myocyte were recorded and analyzed for each condition. All experiments were performed in a closed chamber at 37°C chamber.

### Metabolite Extraction for LC-MS

For perfusate samples, 5uL of perfusate was added to 95uL of 2:2:1 methanol/acetonitrile/water and allowed to incubate on ice for 10 minutes, followed by centrifugation at 17,000g for 15 minutes at 4°C. The supernatant was transferred to autosampler vials with 200uL inserts 4°C until run. For human myocardial tissue, frozen samples were first pulverized with a CryoMill (Restch). The resulting tissue powder was extracted with 2:2:1 methanol:acetonitrile:water (40 uL extraction solvent per 1 mg tissue) overnight at −20°C. The samples were vortexed, and the suspension was centrifuged at 17,000g for 15 minutes. The supernatant was transferred to a clean Eppendorf and allowed to incubate on ice for 15 minutes, prior to a second centrifugation. Supernatant was then added to autosampler vials with 200uL inserts at 4°C until run.

### High Resolution Mass Spectrometry

Mass spectrometry was performed as described previously.^36^ Briefly, a quadrupole orbitrap mass spectrometer (Q Exactive, Thermo Fisher Scientific) coupled to a Vanquish UHPLC System (Thermo Fisher Scientific) with electrospray ionization was set to a scan range of 60 to 550 *m/z* at 1 Hz, with a resolution of 140,000. LC separation was performed on XBridge BEH Amide Column (2.1 × 150 mm, 2.5 um particle size, and 130 Å pore size; Waters Corporation). A gradient of solvent A (95:5 water:acetonitrile with 20 mM ammonium acetate and 20 mM ammonium hydroxide, pH 9.45) and solvent B (acetonitrile) was used for metabolite separation. The flow rate was set to 300 uL/min. The LC gradient was: 0 min, 95% B; min, 95% B; 9 min, 40% B; 11 min, 40% B; 11.1 min, 95% 20 min, 95% B. Mass spectrometry was performed in negative mode for perfusate, and both negative and positive mode for tissues. For untargeted metabolomics the above method was used with the exception of an increased mass range of 70-1000 *m/z*.

### Acyl-CoA extraction and analysis

For quantification and isotope tracing of acyl-CoAs we performed liquid chromatography-high resolution mass spectrometry as previously described in detail (PMID: 26968563). Briefly, for quantification, samples were spiked with 0.1 mL of ^13^C_3_^15^N_1_-acyl-CoA internal standard prepared as previously described from yeast (PMID: 25572876) and 0.9 mL of 10% (w/v) trichloroacetic acid in water. Calibration curves were prepared from commercially available acyl-CoA standards (Sigma Aldrich). Calibration curve samples were also subjected to sonication and extraction in the same manner as the experimental samples. For stable isotope label infused samples, because we included ^13^C_3_^15^N_1_-pantothenate tracing, we used label free quantification based on the peak area or peak area normalized by the weight of the tissue. For isotope tracing of adherent cells, media was aspirated, and cells were quenched in 1 mL 10% (w/v) trichloroacetic acid in water, scraped, and extracted. Samples were homogenized with a probe tip sonicator in 0.5 second pulses 30 times then centrifuged at 17,000 x g for 10 min at 4°C. Then, samples were purified with solid phase extraction (SPE) cartridges (Oasis HLB 10 mg, Waters) conditioned with 1 mL of methanol and then 1 mL of water. Acid-extracted supernatants were loaded onto the cartridges and washed with 1 mL of water. Acyl-CoAs were eluted with 1 mL of 25 mM ammonium acetate in methanol and evaporated to dryness under nitrogen. Samples were resuspended in 50 μL of 5% (w/v) 5-Sulfosalicyilic acid and 10 μL injections were analyzed on an Ultimate 3000 UHPLC using a Waters HSS T3 2.1×100mm 3.5 μm column coupled to a Q Exactive Plus. The analysts were blinded to sample identity during processing and quantification. Data was analyzed using Tracefinder 5.1 (Thermo) and isotopic normalization via FluxFix (PMID: 27887574).

### Cellular Thermal Shift Assay

CETSA was performed as described previously.^37,38^ Briefly, HEK293 or HepG2 cells were cultured in complete DMEM (DMEM containing 10% fetal bovine serum and 100 U/mL penicillin-streptomycin). Separate 20 cm dishes of cells were used for each condition. Cells were treated with EMPA (10 uM), PZ-2891 (10 uM), or Vehicle (DMSO, 0.1%) and allowed to incubate at 37°C for 1.5 h. After incubation, cells were trypsinized, pelleted, and washed with PBS containing cOmplete mini EDTA-free protease inhibitor (Roche). Cells were counted and suspended in PBS to a concentration of 20×10^6^ cells/mL. 100uL of cells were aliquoted into PCR tubes. Cells were subjected to a thermal challenge in a pre-heated Bio-Rad thermocycler from 37°C to 60°C with ∼2.5°C increments for 3 minutes. Cell were removed from heat and allowed to rest at room temperature for 3 minutes before snap freezing in liquid nitrogen. Cells were subjected to 4 freeze-thaw cycles between liquid nitrogen to a 37°C bath. Lysate was collected and centrifuged at 17000g for 40 minutes to precipitate denatured proteins. Supernatant was carefully collected as to not disturb the pellet, mixed with laemelli loading dye and beta-mercaptoethanol (BME) and analyzed by western blot.

### Pank1 Expression and Purification

Pantothenate kinase 1 (Pank1) plasmids were acquired from Addgene (p. 32871) and expanded in lysogeny broth overnight to an optical density of 0.8, after which chloramphenicol (170ug/mL) was added to the culture media and grown for an additional 12 hours. Plasmids were purified by Qiagen miniprep using manufacturer protocol. BL21 complement *E*.*coli* (NEB) were transformed as per manufacturers protocol and selected on agar plates with kanamycin. A single colony was picked and cultured in terrific broth to an optical density of 0.8, at which time the culture was cooled to 4°C and induced by the addition of IPTG (1mM final concentration) and allowed to grown overnight at 18°C with rocking at 220rpm. Bacteria were lysed in HIS-tag wash buffer (300mM NaCl, 10mM imidazole, 50mM NaHPO4, cOmplete mini EDTA-free protease inhibitor) by sonication. The lysate was clarified by centrifugation at 3000rpm for 15 minutes. Supernatant was incubated with HIS-affinity resin (BioRad) for 20 minutes with shaking at 4°C, then washed 5x with HIS-wash buffer. Recombinant protein was eluted with HIS-elution buffer (300mM NaCl, 250mM imidazole, 50mM NaHPO4) and dialyzed with 4-exchanges of kinase storage buffer (50mM Tris-HCl, 5mM dithiothriol (DTT), 10mM MgCl2). Protein was incubated for 10-minutes with phosphate binding resin (Abcam), centrifuged briefly, and stored at −20°C until use.

### Pank1 Kinetic Assays

For malachite green kinase assays: Recombinant HIS-Pank1 was diluted to 20ng/uL in kinase assay buffer (0.1mM Tris-HCl, 10mM MgCl2, 10% Glycerol, pH 7.4). To each well of a 96-well plate was added 50uL of kinase assay buffer containing 145uM ATP and 145uM calcium pantothenate. To respective wells was a concentration gradient of empagliflozin (1.5nM-100uM) or vehicle (DMSO, 0.625% final concentration). To initiate the reaction, 20uL (400ng) of Pank1 was added to each well. Positive controls (PZ-2891) and negative controls (absence of Pank1) were added to each plate. The reaction was allowed to incubate at room temperature for 60-minutes with shaking at 150rpm. Then, 20uL of malachite green reagent (Sigma) was added to each well and allowed to incubate for 30-minutes at room temperature with shaking at 150rpm. Absorbance at 620nM was read using a BioTek Synergy Neo2 plate reader.

### Synthesis of Empagliflozin-Immobilized Resin

Solid-phase resins with immobilized empagliflozin were synthesized utilizing Merrifield-agarose or Epoxy-activated agarose resins. First, the resins (∼1g) were swollen with dichloromethane (DCM) for 1 hour at room temperature. The solvent was then evacuated *in vacuo*, and the resins were washed 3 x w/ 5mL of dimethylformamide (DMF). To a solution of empagliflozin (10eq) in DMF (5mL) was added sodium hydride (60% in mineral oil, 1.2 eq) and allowed to stir at room temperature for 1 minute. The solution was then added to the resins, and the reaction was allowed to proceed for 16h with mixing. The solvent was then evacuated and quenched and the remaining reagent was quenched by the addition of 20% H20:DMF 3x in excess. The resins were then washed 3x with excess DMF. Then, 10eq of ethanolamine in 5mL of DMF is added to the resins, and allowed to incubate for 16h with stirring to fully cap the resins. The resins are then washed 3x with DMF, then 5x with excess PBS to remove organic solvent. The beads were then stored at 4C until use. Hydroxylated-control resins are generated by incubating resins with 10eq of ethanolamine in DMF, and washing as described above.

### Immobilized-EMPA Resin Pull-down

The resins are first suspended in 1mL of PBS to form a slurry. Then, 100uL of resin-slurry is transferred to a 1.5mL Eppendorf tube. To the slurry, is added 100uL of HEK293 cell lysate in M-PER. The resins are then allowed to incubate for 16h at 4°C and with rotating. After incubation, the resins are centrifuged for 1 minute at 10xg, and the supernatant is aspirated. The resins are washed 5x with 1mL of PBS with vortexing, and the resins are allowed to settle to the bottom of the Eppendorf tube by gravity. After washing, protein extraction is performed by adding 100uL of thiourea extraction buffer to the resins and allowed to incubate at room temperature for 30 minutes. The resins are then vortexed, and centrifuged at 10xg for 15 minute. The supernatant is collected, and diluted in 50% glycerol/water with methylene blue for western blot analysis.

### Pank1 – EMPA Co-immunoprecipitation

HEK293 cells grown in complete DMEM with 10% FBS and 1% penicillin-streptomycin are were seeded in 6-well plates and grown to 80-90% confluency. The cells are then washed 1x with PBS, and incubated in DMEM containing 1uM EMPA for 2 hours. After the incubation period, the cells are rapidly washed 3x with ice-cold PBS, and lysed with 200uL of M-PER with protease inhibitor as per manufacturers protocol. The cell-lysate is then clarified by centrifugation at 16xg for 15 minutes, and the supernatant is collected. To clean Eppendorf tubes, 50uL of cell lysate is added followed by 1uL of rabbit anti-PANK1 antibody (CST D2W3N) and allowed to incubate for 16h at 4°C. Controls were incubated in-parallel and did not receive primary antibody. After incubation, magnetic anti-rabbit dynabeads were added (5uL) to each Eppendorf tube. Magnetic beads were incubated for 16 hours at 4°C. The dynabeads were then pelleted through magnification, and the supernatant was aspirated. The beads were then washed 5x with 1mL of PBS with vortexing. Finally, 100uL of LCMS grade water was added to the beads, and boiled at 95°C for 10 minutes. The beads were pelleted by centrifugation 10xg for 10 minutes, and the supernatant was analyzed by HRMS to detect empagliflozin.

### NRVM Isolation and Culture

Neonatal rat ventricular myocytes were isolated as per Pierce-Primary Cell Isolation protocol. Briefly, hearts from p0-2 sprague-dawley pups were harvested and ventricles were isolated from the atria. Ventricles were minced to 1-2mm pieces and digested as protocol states. Cells were separated from debris through a 70 micron-cell filter and pre-plated at 37°C for 90-minutes. Then, cardiomyocytes were harvested by gentle aspiration, centrifuged at 400xg for 5 minutes, resuspended, and plated at a density of 130,000-140,000 cells/well in 12- or 24-well plates, respectively.

### NRVM Tracing Experiments

For CoA tracing in NRVMs, 12-well plates containing 500,000 cells/well were treated with DMEM for primary cells containing stable isotope metabolites (1.3mM 3-C13 Sodium Lactate, 5mM 6,6-D2-Glucose, 0.8mM C13 Valine (Pan), 0.3mM 3-HB, 1mM Glutamine, Final concentrations). The cells were washed 1x with warm PBS, and 1mL of tracing medium was added to each well. Cells were then allowed to incubate at 37°C for 180 minutes. Immediately before harvest, media was aspirated and cells were washed 1x with PBS, and 10% trichloroacetic acid (aq), swirled and transferred to a 1.5mL Eppendorf tube and frozen until analysis.

### Simulation Dataset

We followed a variant of the PopShift protocol to find plausible ligand poses for the Empagliflozin and Acetyl CoA (cite PopShift). To encapsulate the many conformations available to Pantothenate Kinase 1, we built simulations starting from PDB ID 3SMP, a PANK1 crystal structure with Acetyl CoA bound.(cite 3SMP) CoA was removed, along with all ordered waters and coordinated ions, prior to simulation setup. Simulations were set up to use the AMBER ff19SB protein force field, as well as the OPC3 water model using Joung and Cheatham ion parameters for monovalent salts. Potassium was added to bring the molarity of the salt up to 150 mM with 20 Angstroms of initial padding. These simulations were equilibrated on the Bowman Lab cluster using OpenMM 8.1,(cite openmm). We then used Folding@Home to collect 1000 clones of the equilibrated system, each of which was simulated independently for a mean length of approximately 5.1 microseconds (approximately 5.1 milliseconds in aggregate). Simulations were performed in the NPT ensemble using the LangevinMiddleIntegrator set to 300K with a collision frequency of 1 per picosecond and a timestep of 4fs supported by hydrogen mass repartitioning with hydrogens set to 2 AMUs (LMI). Pressure was maintained at 1 atmosphere using the monte carlo barostat (MCBarostat). Frames were saved every 100 ps.

### Markov State Model

We built a Markov State Model (MSM) of the pocket by first selecting any residue within 10 Angstroms of either Acetyl CoA. Because the enzyme is an obligate homodimer, we further included any residue on either chain if it was within this distance on the other chain, totaling 284 residues. We clustered this data the k-hypbrid algorithm as implemented in enspara. Briefly, this consists of k-centers, followed by five iterations of k-medoids, with the RMSD between atomic positions of the superimposed pocket heavy atoms as the clustering metric. 200 centers were used, as this did not result in any ergodic trimming during model building, whereas finer models did. We fit this data using the Bayesian MSM class from Deeptime, with a 50 ns lagtime using the ‘effective’ counts mode to estimate transition counts. From 50 samples from the posterior for the stationary distribution we took the average to obtain equilibrium probabilities.

### Docking

We used GNINA to dock the ligands to the center from each state found during clustering. We aligned each of these structures onto the pocket residues from 3SMP. We drew two boxes, one surrounding each active site, with side-lengths 17×28×28 centered on the center of mass of the Acetyl CoA. GNINA was run with exhaustiveness set to 128, and with the number of poses requested set to 20, in rescore mode. Poses were ranked by their predicted CNN Affinity, and the highest affinity pose was used for subsequent analyses and figure rendering. Docking and score extraction were orchestrated using scripts from the PopShift package.

### Structure analysis

We inspected the state with the most favorable predicted affinity using PyMol (PyMOL). We computed single site affinities using the formulas from PopShift.

Popshift: https://pubs.acs.org/doi/full/10.1021/acs.jctc.3c00870#

3SMP: https://doi.org/10.2210/pdb3SMP/pdb

FF19SB: https://pubs.acs.org/doi/10.1021/acs.jctc.9b00591

enspara: https://pubs.aip.org/aip/jcp/article/150/4/044108/1062329/Enspara-Modeling-molecular-ensembles-with-scalable

OpenMM: https://journals.plos.org/ploscompbiol/article?id=10.1371/journal.pcbi.1005659

Folding@home: https://www.cell.com/biophysj/fulltext/S0006-3495(23)00201-1

LMI: https://pubs.acs.org/doi/10.1021/acs.jpca.9b02771

MCBarostat: https://www.sciencedirect.com/science/article/pii/S0009261403021687

Deeptime: https://dx.doi.org/10.1088/2632-2153/ac3de0

BayesianMSM: https://arxiv.org/abs/1507.05990

PyMOL: https://pymol.org/support.html version 2.5.6

## Data availability statement

All data supporting the findings of this study are available within the paper and its Supplementary Information.

## Acknowledgements

The work was supported in part by a DreamTeam grant from the Penn Cardiovascular Institute and the Children’s Hospital of Philadelphia Frontier Program. NF was supported by the Sarnoff Foundation. ZA was supported by the NIH (HL152446). JJE was supported by the NIH (K08 HL159311). The procurement of human heart tissue was enabled by grants from NIH (R01 HL149891) and the Leducq Foundation to KBM. We would like to thank Gift of Life Donor Program of Philadelphia for enabling the procurement of human hearts from deceased organ donors.

