## Supplemental File for "SGLT2 inhibitors activate pantothenate kinase in the human heart"

**Supplementary Table**

Human Patient Characteristics

| Sample | Etiology | Sex | Age | LV Mass Index | BMI | LVEF | Experiment |
|---|---|---|---|---|---|---|---|
| 1 | NICM | M | 49 | 125.74 | 23.03 | 17.5 | Figure 1 |
| 2 | HCM | M | 31 | 116.82 | 30.46 | 15 | Figure 1 |
| 3 | ICM | F | 44 | 207.32 | 31.96 | 30 | Figure 1 |
| 4 | NICM | M | 52 | 150.38 | 32.82 | 15 | Figure 1 |
| 5 | NICM | F | 46 | 97.98 | 22.03 | 10 | Figure 1 |
| 6 | NICM | F | 26 | 77.66 | 22.58 | 15 | Figure 1 |
| 7 | Sarcoid | M | 61 | 125 | 29.06 | 20 | Figure 1, Figure 2b |
| 8 | NICM | F | 68 | 122.23 | 24.44 | 27.5 | Figure 1 |
| 9 | HFpEF | M | 62 | 106.48 | 25.14 | 57.5 | Figure 1 |
| 10 | ARCV | F | 61 | 127.99 | 24.09 | 70 | Figure 1 |
| 11 | NICM | M | 59 | 147.97 | 32.77 | 10 | Figure 1, Figure 2b |
| 12 | HFpEF | M | 61 | 133.18 | 34.34 | 55 | Figure 2b |
| 13 | HFpEF | M | 58 | 117.79 | 22.40 | 55 | Figure 2b |
| 14 | Sarcoid | M | 59 | 139.33 | 37.22 | 30 | Figure 2b |
| 15 | Congenital | F | 7 | | 19.13 | | Figure 2b |
| 16 | NF | F | 56 | 111.38 | 25.22 | 55 | Figure 2c |
| 17 | NF | F | 56 | 98.39 | 20.70 | 70 | Figure 2c |
| 18 | NF | F | 69 | 139.48 | 33.50 | 52.5 | Figure 2c |
| 19 | NF | F | 61 | 116.12 | 30.43 | 57 | Figure 2c |
| 20 | NF | F | 54 | 114.71 | 36.11 | 55 | Figure 2c |
| 21 | NF | M | 62 | 1411.62 | 23.66 | 65 | Figure 2c |
| 22 | NF | M | 58 | 77.33 | 23.66 | 65 | Figure 2c |
| 23 | NF | M | 20 | 141.62 | 23.66 | 65 | Figure 2c |
| 24 | NICM | F | 46 | 97.98 | 24.03 | 10 | Figure 2c |
| 25 | NICM | M | 28 | 169 | 30.52 | 12 | Figure 2c |
| 26 | Sarcoid | M | 61 | 125 | 29.06 | 20 | Figure 2c |
| 27 | NICM | M | 49 | 125.74 | 23.03 | 17.5 | Figure 2c |
| 28 | HCM | M | 59 | 116.82 | 30.48 | 15 | Figure 2c |
| 29 | NICM | F | 26 | 77.66 | 22.58 | 15 | Figure 2c |
| 30 | NICM | M | 59 | 147.92 | 32.77 | 10 | Figure 2c |
| 31 | NICM | F | 68 | 12.23 | 24.44 | 27.5 | Figure 2c |
| 32 | NICM | M | 52 | 150.35 | 32.82 | 15 | Figure 2c |
| 33 | ICM | F | 53 | 106.59 | 28.52 | 59 | Figure 2c |
| 34 | Sarcoid | M | 59 | 139.33 | 37.22 | 30 | Figure 2c |

**Supplementary Figures**

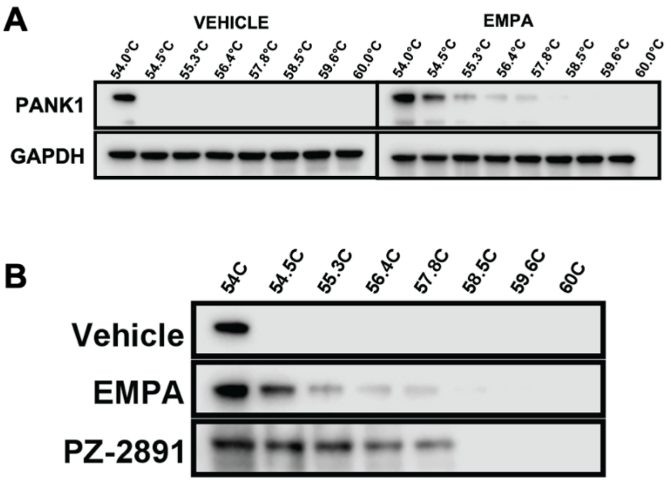


**Supplementary figure 1**. **A,** CETSA of PANK1 from HepG2 cells treated with 10uM Empagliflozin. **B**, CETSA of PANK1 from HEK293 cells treated with vehicle (0.01% DMSO), 10uM Empagliflozin, or 10uM PZ-2891.

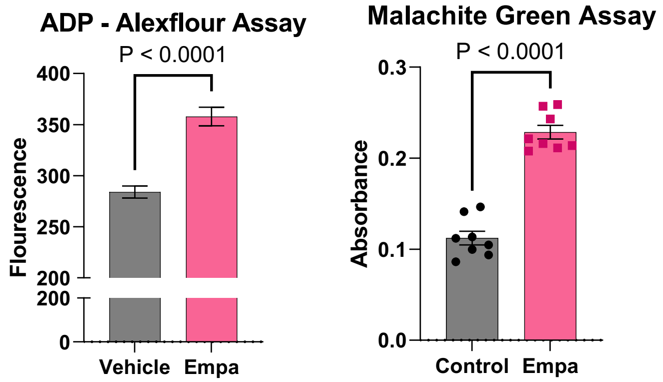


**Supplementary Figure 2**. Single-point enzyme activity assays of PANK1 utilizing 2 different kinase activity assays with 1uM of EMPA *in vitro*. Left: Transcreener ADP Assay (Bellbrook labs) Kit single point kinase activity assay.[1] Right: Malachite green-coupled ADP detection single point kinase activity assay.

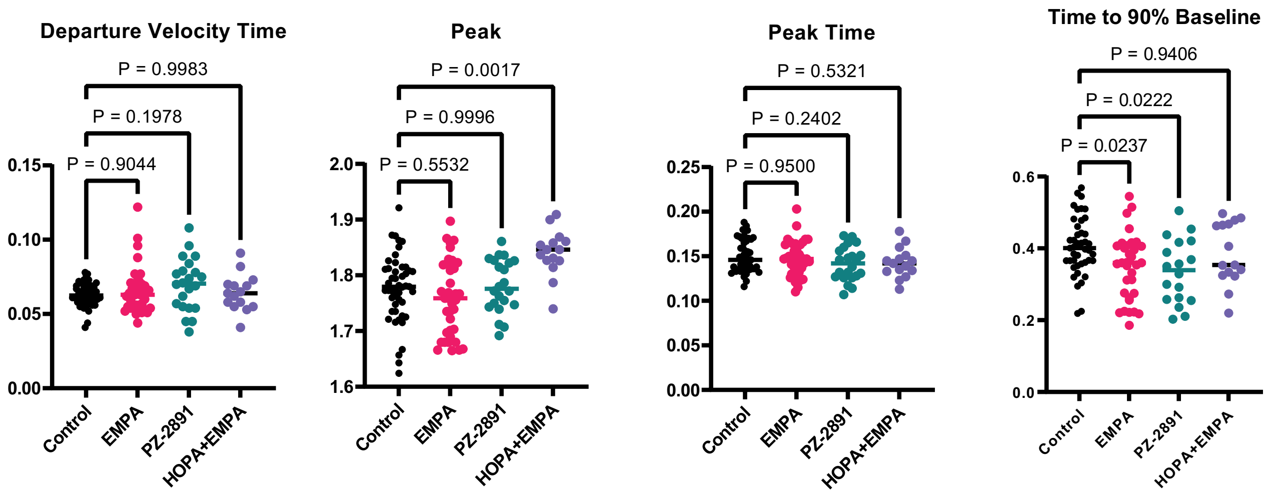


**Supplementary Figure 3**. Additional contractility parameters, accompanying Figure 4.
